# Natural selection has preserved and enhanced the phase-separation properties of FUS during 160 million years of mammalian evolution

**DOI:** 10.1101/2020.03.18.997338

**Authors:** Pouria Dasmeh, Andreas Wagner

## Abstract

Protein phase separation is essential for the self-assembly of non-membraneous organelles. However, we know little about its ability to change in evolution. Here we studied the evolution of the mammalian RNA binding protein FUS, a protein whose prion-like domain (PLD) is essential for the formation of stress granules through liquid-liquid phase separation. Although the prion-like domain evolves three times as rapidly as the remainder of FUS, it harbors absolutely conserved tyrosine residues that are crucial for phase separation. Ancestral reconstruction shows that the phosphorylation sites within the PLD are subject to stabilizing selection. They toggle among a small number of amino acid states. One exception to this pattern are primates, where the number of such phosphosites has increased through positive selection. In addition, we find frequent glutamine to proline changes that help maintain the unstructured state of FUS that is necessary for phase separation. In summary, natural selection has stabilized the liquid-forming potential of FUS and minimized the propensity of cytotoxic liquid-to-solid phase transitions during 160 million years of mammalian evolution.

## Body

The separation of a mixed protein solution into distinct phases is essential for the formation of membrane-less organelles in living cells^1,2^. These biomolecular condensates help mitigate the effects of various stressors, such as energy depletion^3^, increased temperature^3,4^, and drop in cellular pH^4^, which trigger their formation^5,6^.

The RNA binding protein Fused in Sarcoma (FUS)^7-10^ is one of the mammalian proteins whose phase separation is well-studied. FUS is predominantly a nuclear protein and regulates the mRNA life cycle at different stages. In addition, FUS directly interacts with Poly-ADP-ribose polymerase and mediates DNA damage response in the cell^11^. The N-terminal prion-like domain of FUS is essential for the self-assembly of FUS into liquid-like and gel-like states^12,13^. FUS requires this self-assembly for its nuclear functions, such as binding to chromatin and recruitment to the sites of DNA damage^14,15^.

The liquid-like state of FUS is exquisitely sensitive to single point mutations. In fact, missense mutations in FUS occur in patients with the neurodegenerative diseases amyotrophic lateral sclerosis (ALS) and frontotemporal lobar degeneration^8^. For instance, a single ALS-associated mutation, G156E, facilitates a liquid-to-solid phase transition of FUS into irreversible aggregates^16^. The importance of FUS in the life of cells, together with the sensitivity of FUS assemblies to point mutations, raises the possibility that natural selection must actively maintain the ability of FUS to form the liquid-droplet state. We thus hypothesized that evolution has preserved the phase separation propensity of FUS, and avoids the pathological liquid-to-solid phase separation in FUS, just like it maintains folding stability and reduces misfolding in proteins with structured domains^17^.

To validate this hypothesis, we first identified 105 mammalian orthologs of FUS, aligned them (Figure 1A), and computed each residue’s sequence entropy, a widely-used measure of sequence divergence (Figure 1B, see Supplementary Methods for a list of sequences). The PLD domain, which is central for FUS phase separation, has the highest sequence entropy of all FUS domains, with a median ∼3-fold higher than that of the other FUS domains (Figure 1C; Wilcoxon rank-sum test, p=6.12 × 10^−12^), and it shows that the PLD domain evolves much faster than the rest of FUS. Nonetheless, tyrosine residues within this domain are fully conserved, which indicates the essential role of these amino acids for phase separation.

**Figure 1.**
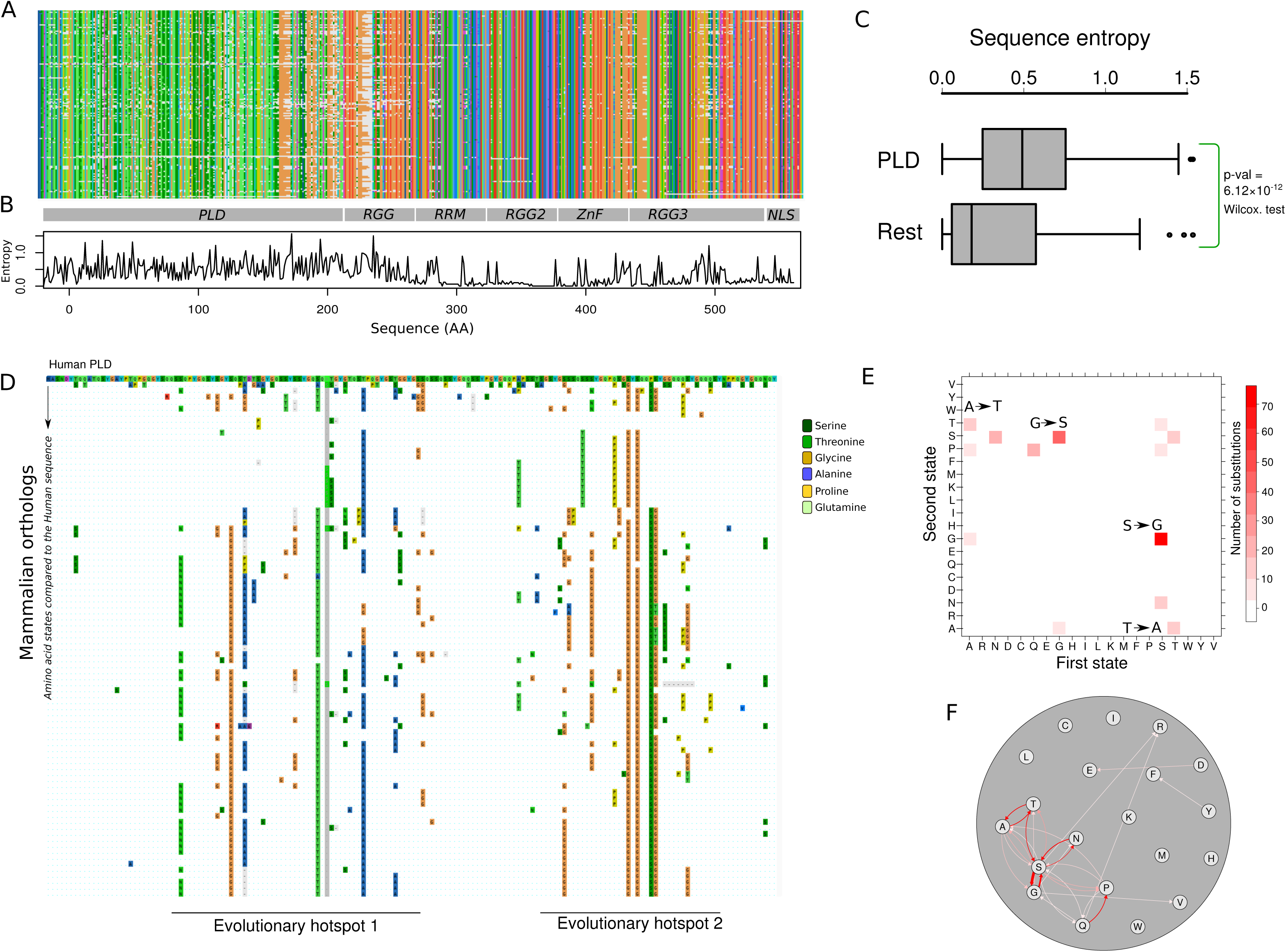
The prion-like domain is the most variable domain of FUS in mammals. A) Multiple sequence alignments of 105 mammalian FUS orthologs colored according to the CLUSTAL color scheme^36^. B) Shannon entropy of each amino acid site in the alignment.C) Box plots comparing the sequence entropy of the amino acid sites in the PLD with the rest of the residues in FUS. D) Substitution map of the PLD in mammals. The first row corresponds to the PLD sequence of human FUS color-coded in the CLUSTAL format. The following rows show the sequences of mammalian PLDs compared to the sequence of the human PLD in the first row. If, at any position, the amino acid is different from that of the human PLD, the new amino acid is shown by colored boxes. Identical amino acids are shown as blue dots. Amino acid substitutions involve changes to serine (forest green), threonine (green), glycine (orange), alanine (blue), proline (yellow), and glutamine (Emerald green). E) Ancestral state mapping of substitutions in the evolution of PLD. Each box represents a substitution with its color saturation proportional to the number of such replacements. The first and second amino acids in a substitution are shown on the X and Y axes, respectively. Arrows highlight the especially frequent A to T, G to S, S to G, and T to A substitutions. F) Ancestral state mapping from panel E represented as a directed graph. Each circle or node represents one amino acid, and substitutions are shown as edges that connect these nodes. The thickness of each edge corresponds to the number of substitutions between the two incident nodes. Substitutions with more than ten occurrences are shown in red.

Within the PLD, we observed two evolutionary hotspots, which are the regions S30 to S86, and A105 to Q147 (All site numbers and amino acids refer to human FUS). These regions are subject to multiple substitutions that involve the amino acids glycine, serine, alanine, threonine, asparagine, proline, and glutamine (Figure 1D). By reconstructing ancestral FUS proteins (Tables S1-S2, see Methods for details), we found that changes where the PLD sites toggle forth and back between G and S (113 changes), as well as between A and T (34 changes), are especially prevalent (Figures 1E-F). Together, these changes account for ∼60% of all changes in the evolution of the PLD. In addition, we found 32 switches between serine and asparagine, and 20 switches between glutamine and proline in these evolutionary hotspots (Tables S1-S2).

To understand whether these amino acid switches are caused by neutral evolution or positive selection, we estimated how strongly evolutionary rates vary across the amino acid sites within the PLD, and along the branches of its phylogenetic tree (see Methods for details)^18^. We detected positive selection in 10 branches (p.val < 0.05) and 12 sites (probability > 0.90) (Figure 2A-B; Table S3) and in three types of substitutions: G to S, S to G and Q to P. We observed the highest likelihood of positive selection for serine at the sites 42, 119, 129, and 131 and threonine at the sites 40 and 71 which occurred in the branches leading to primates and greater apes (Supplementary information).

**Figure 2.**
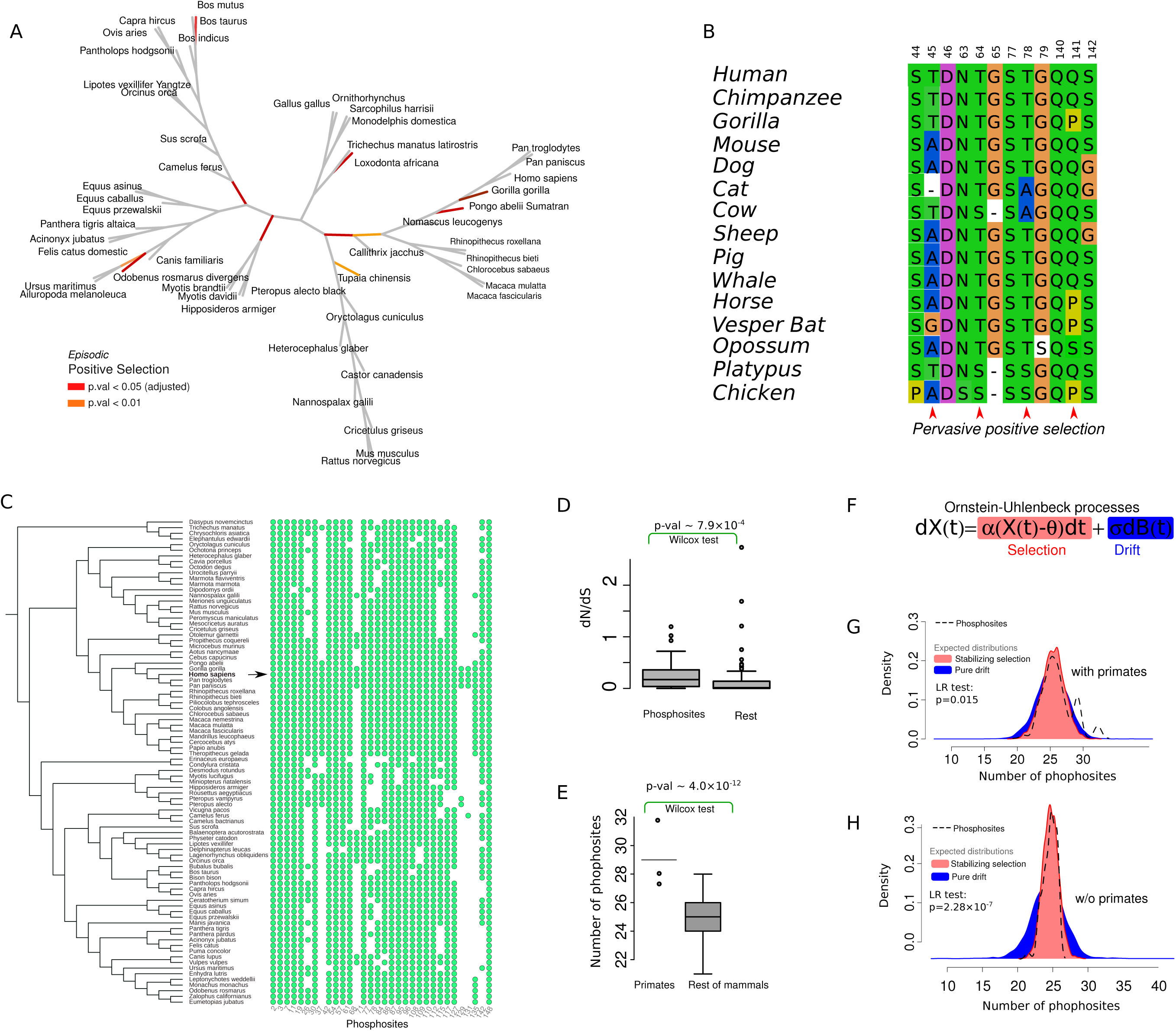
Positive selection in the evolution of phosphorylation sites. A) Mammalian phylogenetic tree showing the branches under positive selection in the evolution of the PLD. Branches on which positive selection is detected with significance levels of p=0.01 and p=5×10^−4^ (adjusted p=0.05, using Bonferroni correction^37^ with 97 branches), respectively, are colored in orange and red (P-values corrected for multiple sampling). B) Sites under pervasive positive selection shown in a subset of mammalian sequences. C) Mammalian phylogenetic tree, along with a map indicating the presence (green circle) or absence (blank) of serines/threonines in the phosphorylation sites of human FUS (arrow). D) Bboxplots comparing the rate of evolution (dN/dS) of phosphosites with the rest of the PLD residues. E) Boxplots comparing the total number of phosphosites in primates with this number in the rest of mammals. F) The general stochastic equation of an Ornstein-Uhlenbeck process with drift and selection components, highlighted in blue and red, respectively. G and H) Probability density of the number of phosphosites (dashed line) compared with the probability density of the evolution of phosphosites under drift (blue) and stabilizing selection (red). The expected blue and red distributions are obtained by simulating the Ornstein-Uhlenbeck process 100 times for all the mammalian PLD sequences (panel G), and for the mammalian PLD sequences without primates (panel H). LR: likelihood ratio.

The positively selected residues in primates (i.e., sites 42, 71, 78, 129, and 131) are among the sites in human FUS that are phosphorylated at serine and threonine^19,20^. Their phosphorylation not only increases the recruitment of FUS to the sites of DNA damage^21^, but also inhibits its liquid-to-solid phase separation^19^. From a total of 32 phosphosites, only nine sites (i.e., sites 3, 7, 11, 26, 57, 77, 87, 96, and 148) were fully conserved, but the rest (24 sites) switched forth and back between only two pairs of amino acids (G-S and A-T) (Figure 2C). These sites occurred in both evolutionary hotspots, and their evolutionary rates were significantly higher than for the rest of the PLD residues (Figure 2D; Wilcoxon rank-sum test, p=7.9×10^−4^). The PLD sequences of primates and great apes harbor an exceptionally large number of phosphosites. That is, they harbor 29 and 31 phosphosites, respectively, which is 3 and 6 sites more than the average number of mammalian PLD phosphosites (Figure 2E). Therefore, positively selected G to S and A to T substitutions have significantly increased the total number of phosphosites in the PLD sequences of primates. (Wilcoxon rank-sum test, p=4.0×10^−12^).

Aside from primates, the average number of phosphosites in the mammalian PLD sequences is ∼ 26 ± 2 sites. This small variation might indicate that the total number of S/T amino acids in these sites are stabilized in the evolution of the FUS in mammals. To find out whether stabilizing selection has acted on our FUS sequences, we compared the likelihood that genetic drift alone or drift together with selection acted on the total number of phosphosites using an Ornstein-Uhlenbeck process ^22^. This process has been used to compare the likelihood of drift alone with that of drift and selection in the evolution of different traits and characters^23^ (Figure 2F; see Methods for details). We found that stabilizing selection better explains the evolution of phosphosites than pure drift (likelihood ratio test, p=0.015, Figure 2G; Table S4). Interestingly, this signature of stabilizing selection increases dramatically when primates are removed from this analysis (Figure 2H; likelihood ratio test, p=2.28×10^−7^; Table S4). Together with our phylogenetic analysis of positive selection (Figures 2A-B), these observations suggest two regimes in the evolution of FUS phosphosites. In mammals except for primates, the number of phosphosites is under stabilizing selection. In primates and, in particular, great apes, positive selection has further increased the number of phosphorylation sites.

The disordered domains of proteins, in particular proteins that undergo phase separation, preserve key amino acid features such as charge and sequence composition throughout their evolution^24,25^. We thus examined the physicochemical properties that are either conserved or positively selected in the evolution of the PLD in mammalian FUS (see Methods for details). We found that amino acid substitutions in the PLD have significantly conserved polarity, flexibility, and solvation free energy (Figure 3A, Table S5; Chi-square goodness-of-fit, p < 10^−7^). We also found several properties whose changes were more frequent than expected from strict neutrality, and that had diversified in the evolution of the PLD (Figure 3A). The most significantly diversified property is the average occurrence of amino acids in a tetrapeptide unit in protein structures^26^. This property quantifies the nucleation propensity of amino acids in segments of four residues, and divides the amino acids into two groups. Tetrapeptides with amino acids in the first group (Pro, Gly, His, Tyr, Cys, Asn and, Trp) are more likely to adopt extended structures. Tetrapeptides with amino acids in the second group (all other amino acids), are more likely to form helical and bend conformations.

**Figure 3.**
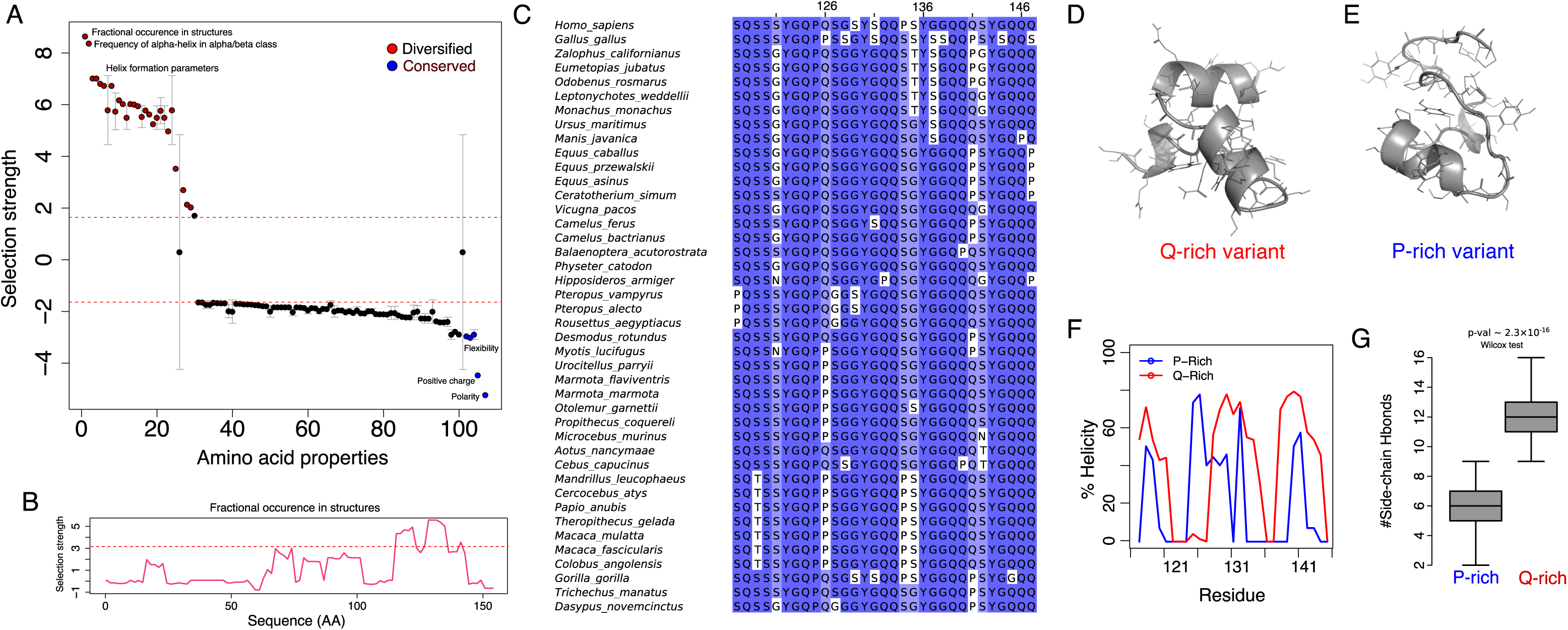
Proline substitutions disrupt the formation of secondary structure in an evolutionary PLD hotspot. A) 108 different amino acid properties showing radical changes in the evolution of the PLD, ranked by selection’s tendency to either help diversify them (positive values, red) or conserve them (negative values, blue) during PLD evolution. Properties falling above and below the dashed line are selected for or against at a high significance level of 0.001. Phosphosites are excluded from this analysis. Because several amino properties correlate with each other, we averaged the selection’s tendency between highly correlated properties (Spearman’s correlation R > 0.8). Error bars show one standard deviation of selection strength between highly correlated amino acid properties. Selection tends to diversify properties that change the structure-formation propensity, while it tends to conserve amino acid polarity and flexibility. B) Selection strength of the property ‘fractional occurrence in tetra-peptides in protein structures (RACS820101^26^)’ over the length of the PLD (horizontal axis). The area highlighted in red shows the region S117 to S147, where selection to diversify this property is maximal. C) Multiple sequence alignment of the segment S117 to S147 for selected mammalian PLD sequences. Color saturation represents the frequency of the amino acid in each column, and ranges from dark blue (> 80%) to white (<40%). D) PEP-FOLD predicted structure of the glutamine-rich, and E) the proline-rich variant of the region S117 to S147. The sites 126, 132, 140, 141, 146, and 147 were computationally substituted to glutamine or proline to create the Q-rich or the P-rich variants, respectively. Both structures were generated by the PEP-FOLD webserver. F) The percentage of simulation time that any one residue (horizontal axis) is found in a helical conformation and G) the number of side-chain hydrogen bonds in the molecular dynamics simulations of the Q-rich (red) and the P-rich (blue) variants. The p-value was calculated with Wilcoxon rank-sum test.

For this property, amino acid substitutions in the region S117 to S147 had the maximum strength of positive selection (Figure 3B, Table S6). This region is enriched in proline substitutions (i.e., in residues 105, 117, 126, 132, 134, 140, 141, 146, and 147; Figure 3C). Importantly, our analysis of positive selection had shown that Q to P substitutions are positively selected in different branches of the phylogenetic tree (e.g., Q141P in Gorilla (*Gorilla gorilla*) and S134P in the branch leading to primates; Supplementary information).

To study the effect of proline substitutions in this evolutionary hotspot, we made a glutamine-rich and a proline-rich variant from the region S117 to S147 by selecting Q and P in all residues that had experienced Q to P substitutions in different mammalian sequences (residues 126, 132, 140, 141, 146, and 147), respectively (Supplementary information). We then predicted the secondary structure content of the two variants using the PEP-FOLD algorithm^27^ and validated the stability of these predictions using molecular dynamics simulations (see Methods for details). The glutamine-rich variant forms three short helices that extend from residue Q118 to S121, from S129 to S135, and from Q139 to G144 (Figure 3D). In contrast, the proline-rich variant is mostly unstructured and retains only partially the middle of the three helices (Q126 to Y129; Figure 3E). Importantly, the Q-rich variant showed higher helical content (Figure 3F) and, on average, five more side-chain hydrogen bonds compared to the P-rich variant (Figure 3G). We thus conclude that Q to P substitutions help maintain an unstructured state that is necessary for self-assembly and phase separation of FUS.

Finally, we examined the changes in the propensity of fibril formation in the evolution of the PLD in mammalian FUS. The first hotspot in the PLD, from S30 to S86, corresponds to a region that forms a fibrillar beta-sheet structure at high concentrations (Figure 4A)^12^. The formation of these cross-beta sheet structures has been proposed^12^, and disputed^28,29^, to drive phase separation of FUS. Strikingly, we observed hydrogen-bond breaking substitutions in these regions that abolish side-chain hydrogen bonding and likely destabilize the fibril core (Figure 4B). For example, T78 and S84, which form inter-residue hydrogen bonds in the structure of the fibril core, are repeatedly substituted to alanine and glycine in different mammals. Other examples include alanine or proline substitutions in the residues S48, Q69, and T71, which hydrogen-bond and join the segment S44 to Y50 with T64 to G80. We further predicted the stability of fibril cores in different mammals and found a substantial variation in free energy of folding (Figure 4C, Table S7) which is more likely caused by pure drift than stabilizing selection (likelihood ratio test, p=0.019; Table S8). In line with this observation, evolutionary rates of the PLD sequences were not significantly higher along the branches leading to fibrils with higher stabilities (Spearman’s rank correlation, R=0.047, p=0.71). Altogether, our analyses reveal that stability of the fibril core is unlikely maintained and selected in the evolution of the PLD in mammalian FUS.

**Figure 4.**
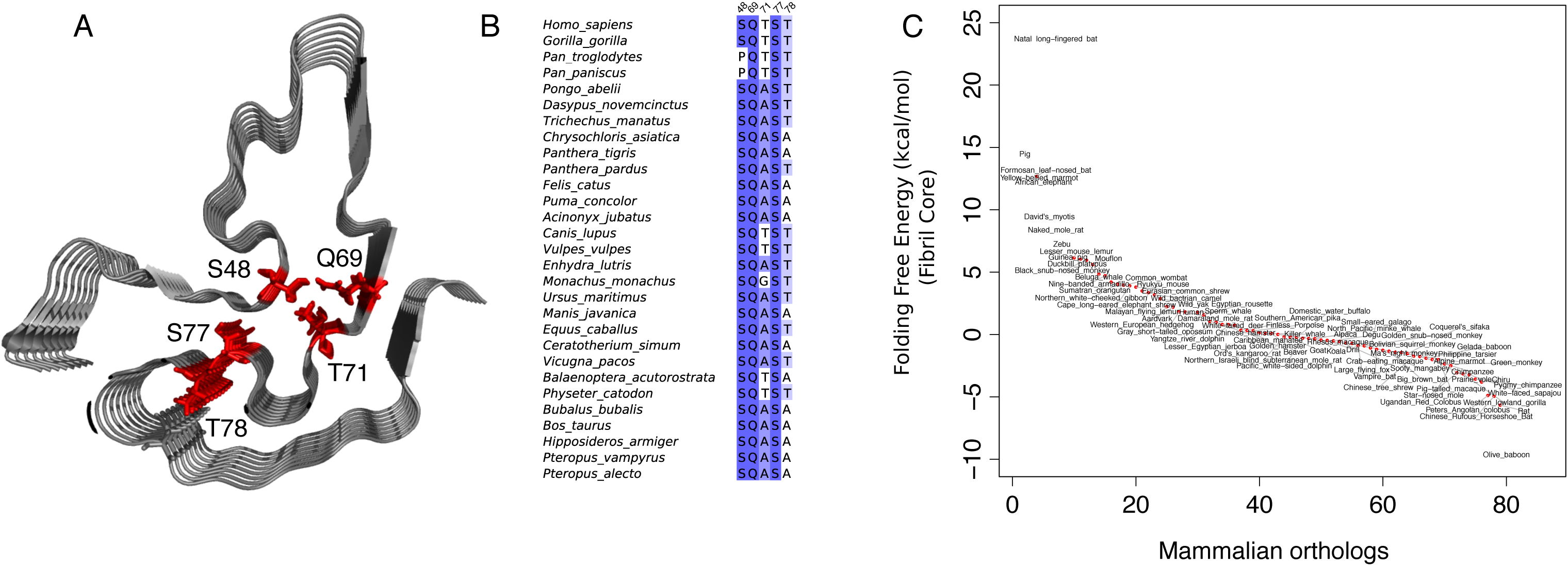
The stability of the fibril core varies substantially among mammalian PLD sequences. A) Structure of the fibril core in the human PLD (PDB ID: 5W3N^11^) with the residues S48, Q69, T71, S77, and T78 shown in red. B) Sequence alignment of these residues for selected mammals. C) Predicted free energy of folding of the fibril core, ranked for different mammalian PLDs. The stability of the human fibril core is taken as the reference free energy level of zero.

In summary, our work shows that properties affecting phase separation are highly evolvable in FUS, an essential protein that is also involved in neurological diseases. The lack of protein structure in the prion-like domain of FUS leads to substantial sequence variation, a feature that is common in intrinsically disordered proteins^30-33^. Given this high overall divergence, stabilizing the condensed liquid droplet state requires evolutionary mechanisms to maintain the disordered nature of the protein and avoid liquid-to-solid phase transitions. Our observations show that stabilizing selection of phosphorylation sites and positive selection of proline substitutions are two primary mechanisms to maintain the phase separation propensity of FUS in mammals. Previously, proline substitutions have been shown to reduce the formation of irreversible aggregates of the huntingtin protein in Huntington disease^34^. Our analysis extends this observation to the much greater evolutionary time scale of mammalian evolution, which unfolded over 160 million years.

FUS is only one among ∼ 2600 proteins in the human proteome whose sequence architecture is similar to FUS, and our observations may apply to many of these proteins as well. These proteins, include the members of the FUS-like family of proteins such as EWS and TAF15, which have prion-like and RNA-binding domains that are similar in length and composition to these domains in the corresponding FUS domains, and might function as scaffolds for biomolecular condensates in the cell^35^.

## Supplementary Information

### 1. Supplementary Figures

**Figure S1.**
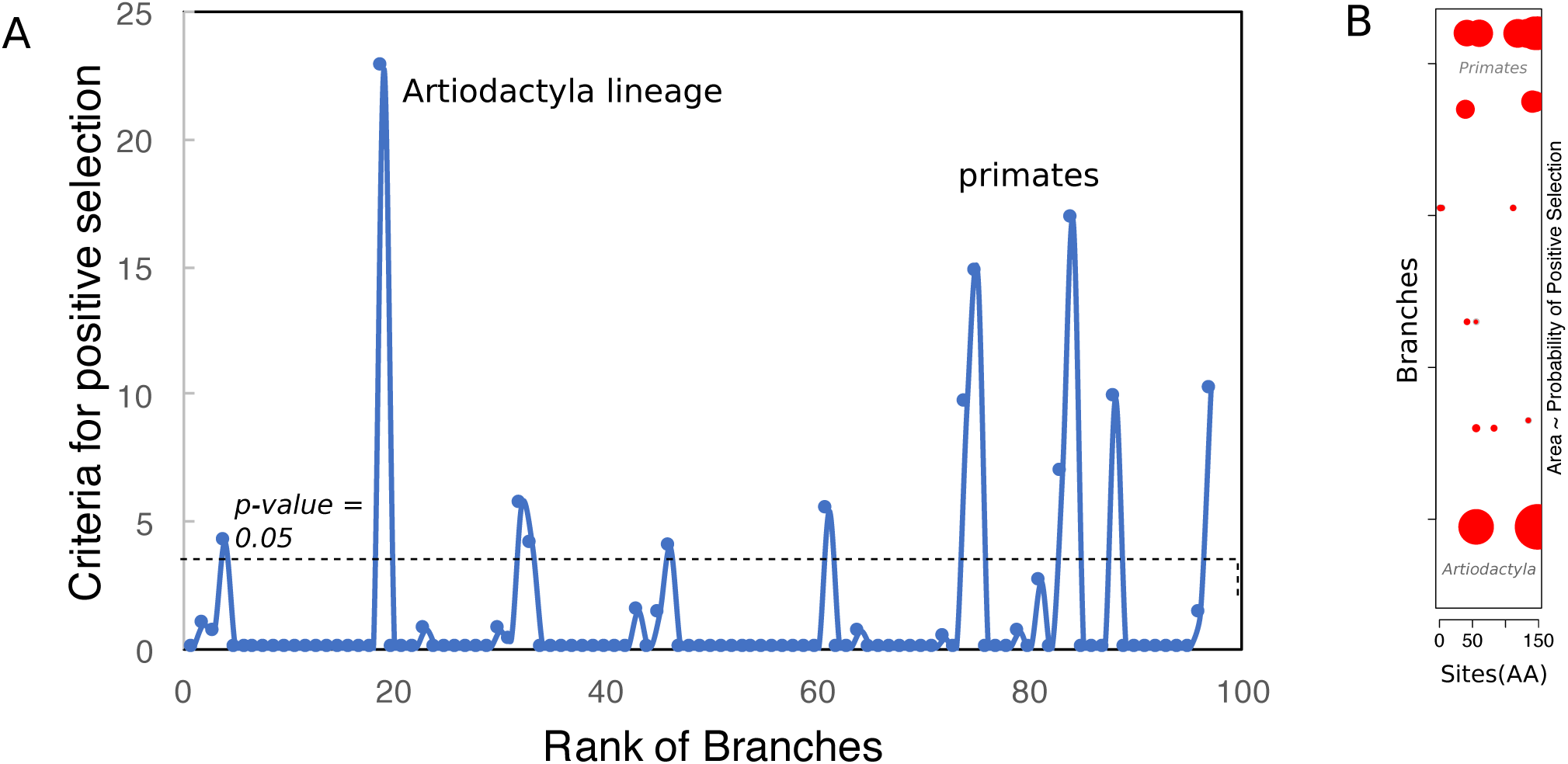
A) The criteria for positive selection (Twice the difference in the logarithm of likelihood function) for different branches of the phylogenetic tree. The branches leading to species in the orders Artiodactyla and primates have significant probability of positive selection. B) The probability of per-branch selection of amino acid sites (i.e., the branch probability multiplied by the site probability) for different branches of the phylogenetic tree.

**Figure S2.**
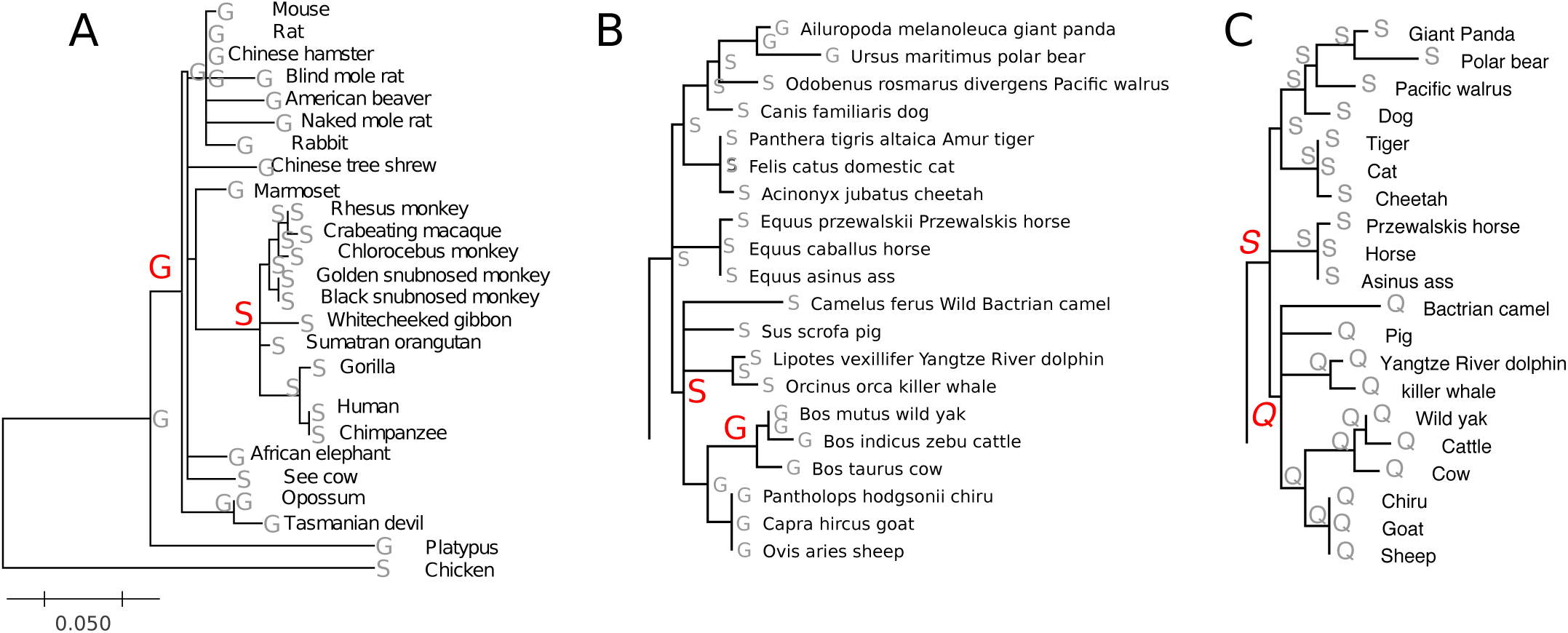
Substitution patterns mapped on the phylogenetic tree of mammals in three representative sites under positive selection, A) S147, B) G56, and C) 148. See the supplementary results for the list of all transitions under positive selection.

**Figure S3.**
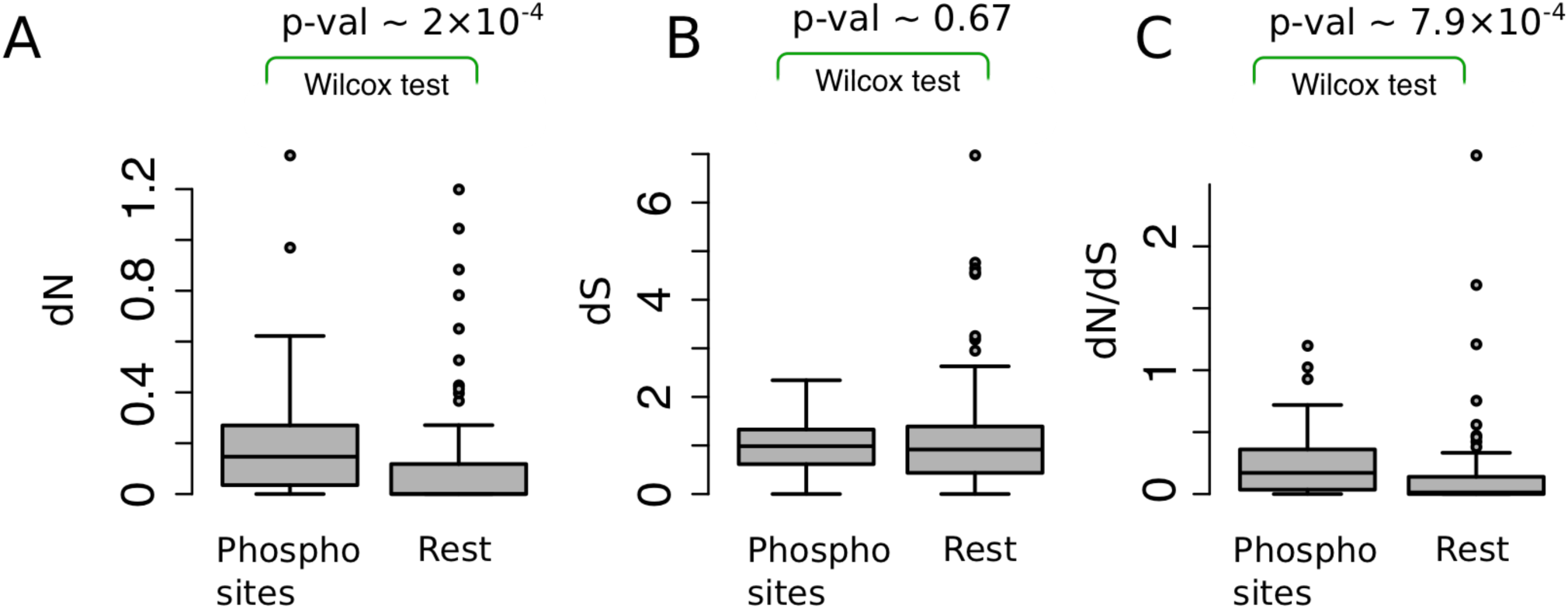
The rate of nonsynonymous substitutions, d*N* (panel A), Synonymous substitutions, d*S* (panel B), and evolutionary rate, d*N*/d*S* (panel C), for phosphorylates sites and the rest of the PLD residues.

**Figure S4.**
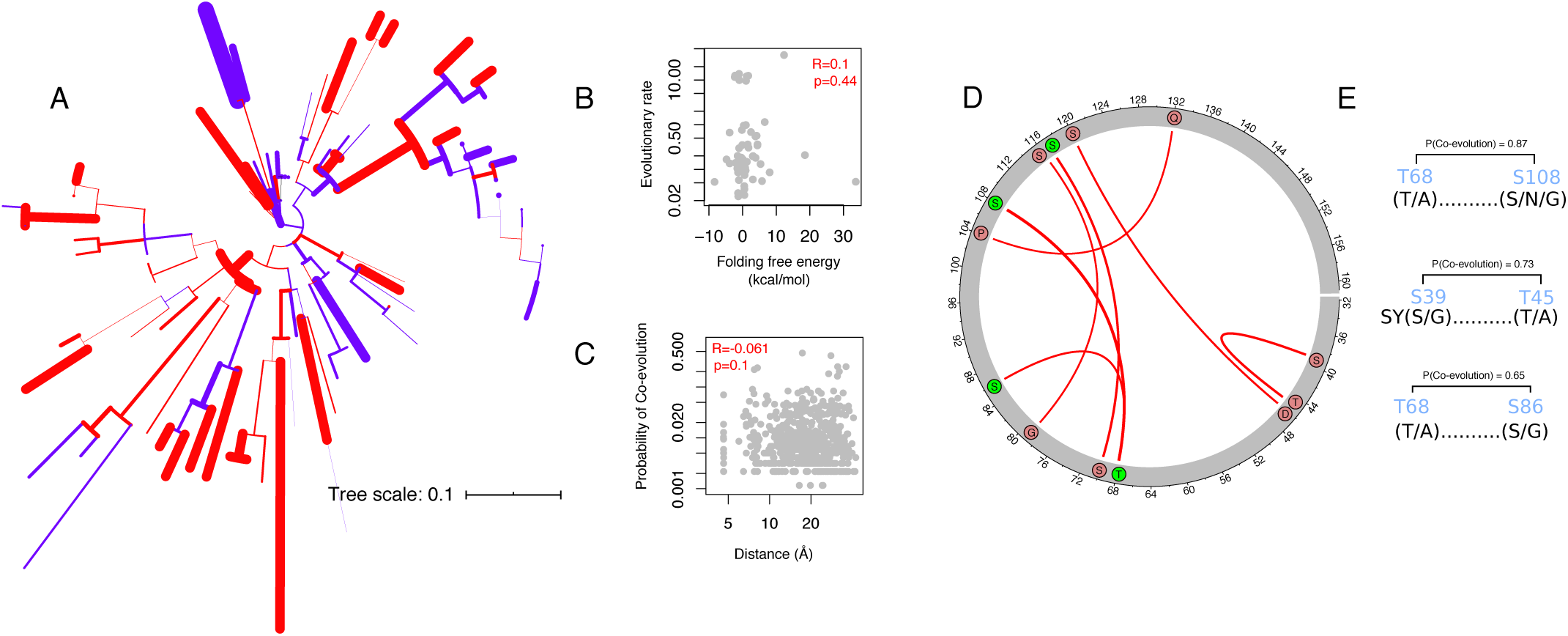
A) Mammalian phylogenetic tree, showing branches with more stable (red) or less stable (blue) fibril cores compared to the structure of the human fibril core. B) Evolutionary rate (d*N*/d*S*) of the PLD in the branches of the phylogenetic tree versus the change in the predicted folding free energy of the fibril core compared to the human fibril core. C) Probability of co-evolution between pairs of residues versus their distance on the PDB structure of the human fibril core (PDB ID: 5W3N^11^). ‘R’ indicates Spearman’s rank correlation coefficient. D) Circos map showing PLD residue pairs that co-evolve with high probability (red, p > 0.5). Residues phosphorylated in human FUS are shown as green circles. E) High probability of co-evolution observed in the pairs T68-S108, S39-T45, and T68-S86. For each residue, the most frequent amino acids are shown in parenthesis.

### 2. Supplementary Tables

- Table S1: The substitution patterns of the ancestrlal reconstruction in the PLD of FUS
- Table S2: The substitution patterns of the ancestrlal reconstruction in the PLD of FUS
- Table S3: The likelihoods for the alternative and the null model of branch-site test for positive selection
- Table S4: Ornstein-Uhlenbeck processes for phosphorylated sites (without primates)
- Table S5: Highly conserved and highly diversified physicochemical properties in the evolution of the PLD in FUS.
- Table S6: Z-score values of the property RACS820112 per sliding window of 5 codons.
- Table S7: FoldX free energy terms for extant and ancestral PLD sequences of mammalian FUS.
- Table S8: Ornstein-Uhlenbeck process for changes of the folding free energy in the evolution of fibril core in FUS.

### 3. Supplementary files

All data are available at: https://figshare.com/articles/_/11993265

- Sequences of mammalian FUS and ancestral sequences.
- Structures of Q-rich and P-rich variants made by PEP-fold algorithm.
- Structure of fibril cores in mammals made by FoldX *in silico* mutagenesis.

### 4. Supplementary Methods

#### Data compilation

We retrieved 105 coding sequences of mammalian FUS genes from the NCBI^1^ and ENSEMBL^2^ databases. We subdivided these sequences into three subsets. We used the first of these subsets, which comprised 105 mammalian sequences, to build a multiple sequence alignment for the calculation of sequence entropy. We used the second subset of 85 sequences, which had confident phylogenetic support from the TimeTree database^3^ to i) estimate the likelihood of drift and selection with the aid of an Ornstein-Uhlenbeck process, ii) correlate the evolutionary rates of the PLD with the stability of the fibril core’s structure, and iii) to analyze co-evolution of the PLD residues. Finally, we used a third subset of 50 sequences with diverse taxonomic sampling in our analysis of positive selection. (The accession numbers of all sequences used in this work, as well as the sequences of reconstructed ancestors are available in the online supplementary information).

#### Estimating evolution rate and detecting positive evolution

We prepared protein sequence alignments with the codon-based CLUSTAL algorithm^4^ implemented in MEGA^5^ and Aliview^6^ using the default parameters. We used the codeml program within the PAML suite^7^ to obtain maximum-likelihood estimates of the ratio d*N*/d*S*, i.e., the ratio of the number of nonsynonymous substitutions per nonsynonymous site to the number of synonymous substitutions per synonymous site. This ratio is a widely-used measure of selection strength on an evolving sequence^8^. For the estimation of d*N*/d*S*, we used the equilibrium codon frequencies from the products of the average observed frequencies in the three codon positions using the F3X4 model^7^. We tested the likelihood of positive selection in our sequences using the branch-site test for positive selection^9^. In this model, a phylogenetic tree is partitioned into the foreground and background branches. The likelihoods of dN/dS>1 and dN/dS=1 along the foreground branches are compared using likelihood ratio tests. We determined the posterior probabilities that specific sites (amino acids) are subject to positive selection using the Bayes Empirical Bayes (BEB)^10^ method implemented in PAML^7^.

#### Ancestral Sequence Reconstruction

To reconstruct ancestral sequences, we fitted different substitution models to our data (PLD sequences and the mammalian phylogenetic tree), allowing that evolutionary rates may vary among protein sites. The substitution model JTT^11^ with the gamma distribution of evolutionary rates had the highest Bayesian Information Criterion (BIC) score^35^. We thus used this model and inferred ancestral PLD sequences using the Maximum Likelihood method implemented in MEGA7^5^.

#### Detection of amino acid properties under selection

We used the TreeSAAP method^12^ to infer how natural selection may change amino acid properties in the evolution of the PLD. Briefly, this method compares the distribution of changes in amino acid properties along the branches of a phylogenetic tree with an expected distribution, using the codon composition of a set of extant sequences. Changes in amino acid properties are divided into eight categories, from the most conserved (category 1) to the most radical changes (category 8). The method then calculates the goodness of fit (*χ*^2^-distribution) between the expected and the observed frequencies and tests the hypothesis that these distributions are equal for each amino acid property. For a specific property, the deviation between observed and expected frequencies in each category is calculated using a Z-score. We refer to this Z-score as the deviation from neutrality or the selection strength throughout this paper. A highly significant z-score (z > 3.09, p<0.01) shows that more non-synonymous substitutions change the property of interest compared to neutral evolution.

#### Detection of Co-evolution

To infer the co-evolutionary history of protein sites within the PLD, we used Bayesian Graphical Models^13^ implemented in the HyPhy package^14^. We first used our ancestral reconstruction of FUS to construct a binary matrix representing the presence and absence of substitutions on each branch (rows) of the phylogenetic tree and in each site of the protein (columns). The joint distribution of all substitutions was then inferred using Bayesian networks and Markov Chain Monte Carlo (MCMC) sampling with default parameters in the SpiderMonkey method^13^. We avoided the use of mutual information to infer co-evolution because it leads to a high rate of false positives in the detection of co-evolving sites when sequences are substantially similar^19^ (∼ > 62%), as in our case.

#### Prediction of folding free energy

We predicted the stability of the structure of the fibril core of the PLD in different mammals. We generated the 3D structures of the fibril core using its structure in human (PDB ID: 5W3N^15^) as a template. We then calculated the free energy of folding of the fibril core made from the mammalian PLD sequences using the FoldX algorithm, which uses an empirical force field for the prediction of the free energy change of protein structures upon mutations^16,17^. We first minimized the free energy of this structure using the Repair command in FoldX^17^. We then created *in silico* mutants of this structure to create different mammalian PLD orthologs using the BuildModel command of FoldX^17^.

#### Ornstein-Uhlenbeck processes

To estimate the significance of stabilizing selection versus pure drift, we used Ornstein-Uhlenbeck processes that are corrected for phylogenetic dependence of species^18^. These models have been used to test various evolutionary hypotheses in the evolution of different characters and traits^19^, gene expression level^20^, and protein structure^21^. In brief, these models assume that the character of interest, X(t), evolves in time unit (t), according to an Ornstein-Uhlenbeck process:

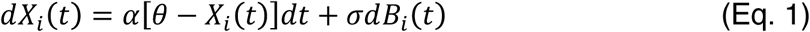

The parameter *θ* is the optimum value of X(t) in the *i*^th^ lineage and the parameters *α* and *σ* represent the strength of selectin and drift, respectively. The term dB_i_(t) is a white noise term, with mean 0 and variance dt. Equation 1 defines a Gaussian process whose moments depend on the parameters *θ, α*, and *σ* and the total time spent for a character in the lineage of interest, t=T. T is either defined as the evolutionary time or equal to the branch length, as in our case. The values of the optimum parameter of the evolving trait (*θ*) can vary according to the assumed evolutionary process. For example, if the character of interest evolves under a single optimum, *θ* is the same for all lineages. These parameters are estimated by minimizing the logarithm of a likelihood function that assumes multivariate normality of all characters at the terminal taxa, X(t=T).

We used the total number of phosphosites and the folding free energy as the traits of interest in evolution. We then fitted the models for pure drift and stabilizing selection using the BROWN and HANSEN commands in the OUCH package^19^, respectively. The input to these commands was the mammalian phylogenetic tree in Newick format, together with a data vector of the trait of interest, either the total number of phosphosites or the stability of the fibril core. We used the initial values of *α* = 1 and *σ* = 1 to initialize the optimization process using the Nelder and Mead simplex algorithm^22^. For modeling stabilizing selection, we assumed that all nodes belong to a single selective regime. We used likelihood ratio tests to compare the likelihoods of drift and selection.

#### Prediction of peptide structure and molecular dynamics simulations

We generated the initial structures of the glutamine-rich and the proline-rich variants using the PEP-FOLD online web server^23^. For molecular dynamics simulations, we used the GROMACS package^24^ (v2019a) and employed periodic boundary conditions at 300 K and 1 atm, with a time step of 2 fs. We chose the Gromos 54a7^25^ force field because of its ability to reproduce the kinetics of helix formation^26^. We kept the temperature and pressure constant with the Nose-Hoover thermostat^27,28^ (time-constant = 0.1 ps) and the Parinello-Rahman barostat^29^ (time constant = 1.5 ps), respectively. For both van der Waals and short-ranged Coulombic interactions, we used a cut-off radius of 1.0 nm and used the particle-particle mesh Ewald method for the long-ranged Coulombic interactions^24^. We minimized the energy of both structures by the steepest descent method, followed by a position-restraint simulation to equilibrate the water molecules. We then performed a grand canonical ensemble simulation (constant number of particles, temperature, and pressure) at 300K for 20 ns and calculated the percentage helicity of different residues and the number of side-chain hydrogen bonds. We performed all statistical analyses in R, using scripts available on GitHub (https://github.com/dasmeh/FUSEVOL).

### 5. Supplementary Results

#### Co-evolution of residues in the PLD

We also pursued a complementary analysis to study the co-evolution of fibril core residues. If the fibril core has remained intact during mammalian evolution, then we would expect that residues in the fibril core co-evolve with each other, and possibly also with the rest of the PLD33. To find out, we used Bayesian Graphical Models^13^ to estimate the probability of co-evolution between pairs of residues in the fibril core (see Methods). Despite the presence of pairs of co-evolving residues (i.e., S39-T45, T68-S86, S68-S108, and G79-S116), we found no correlation between the probability of co-evolution and the distance of co-evolving residues on the 3D structure (Spearman’s rank correlation, R=- 0.061, p=0.13). Interestingly, the co-evolving residues T68 and S86 are phosphorylated in human FUS, showing that co-evolution has modulated the phosphorylation potential of the PLD. Altogether, our analyses reveal that stability and the structural integrity of the fibril core are unlikely maintained in the evolution of the PLD in mammalian FUS.

#### Positive selection patterns along the branches

1. Branch Number: 4 (Site, Probability, Transition) 64, 0.674, T > S
2. Branch Number: 19 (Site, Probability, Transition) 56, 0.776, S>G 148, 0.955, S>Q Lineage in the phylogeny: ((((((((((((Bos_mutus_wild_yak:0.019457,Bos_indicus_zebu_cattle:0.006900):0.006263,Bos_taurus_cow:0.013914):0.070474,(Pant holops_hodgsonii_chiru:0.018526,(Capra_hircus_goat:0.000004,Ovis_aries_sheep:0.006562):0.014510):0.045189):0.147414,(Lipot es_vexillifer_Yangtze_River_dolphin:0.025717,Orcinus_orca_killer_whale:0.027657):0.054548):0.026334,Sus_scrofa_pig:0.110769):0.000969,Camelus_ferus_Wild_Bactrian_camel:0.169220):0.089605 ←
3. Branch Number: 32 (Site, Probability, Transition) 56, 0.728, S>G 83, 0.617, S>G Lineage in the phylogeny: (((Ursus_maritimus_polar_bear:0.043702, Ailuropoda_melanoleuca_giant_panda:0.009459):0.034020 ←
4. Branch Number: 33 (Site, Probability, Transition) 135, 0.690, G>T Lineage in the phylogeny: Odobenus_rosmarus_divergens_Pacific_walrus:0.050105 ←
5. Branch Number: 46 (Site, Probability, Transition) 42, 0.819, S>G 56, 0.604, S>G Lineage in the phylogeny: ((((((((Bos_mutus_wild_yak:0.019457,Bos_indicus_zebu_cattle:0.006900):0.006263,Bos_taurus_cow:0.013914):0.070474,(Panthol ops_hodgsonii_chiru:0.018526,(Capra_hircus_goat:0.000004,Ovis_aries_sheep:0.006562):0.014510):0.045189):0.147414,(Lipotes _vexillifer_Yangtze_River_dolphin:0.025717,Orcinus_orca_killer_whale:0.027657):0.054548):0.026334,Sus_scrofa_pig:0.110769):0.000969,Camelus_ferus_Wild_Bactrian_camel:0.169220):0.089605,(((Equus_asinus_ass:0.006650,Equus_caballus_horse:0.00000 4):0.000004,Equus_przewalskii_Przewalskis_horse:0.006681):0.122260,((Panthera_tigris_altaica_Amur_tiger:0.000004,(Acinonyx_j ubatus_cheetah:0.006612,Felis_catus_domestic_cat:0.013246):0.000004):0.052309,(((Ursus_maritimus_polar_bear:0.043702,Ailur opoda_melanoleuca_giant_panda:0.009459):0.034020,Odobenus_rosmarus_divergens_Pacific_walrus:0.050105):0.016635,Canis_ familiaris_dog:0.040800):0.021089):0.042717):0.000004):0.012372,((Myotis_brandtii_Brandts_bat:0.071429,Myotis_davidii_Vesper _bat:0.034392):0.211993,(Hipposideros_armiger_great_roundleaf_bat:0.227675,Pteropus_alecto_black_fruit_bat:0.067987):0.0446 56):0.044726 ←)
6. Branch Number: 61 (Site, Probability, Transition) 112, 0.598, S>G (Tupaia_chinensis_Chinese_tree_shrew:0.168529 ←)
7. Branch Number: 74 (Site, Probability, Transition) 40, 0.998, G>T Lineage in the phylogeny: (Pongo_abelii_Sumatran_orangutan:0.021484 ←)
8. Branch Number: 75 (Site, Probability, Transition) 141, 0.761, Q>P Lineage in the phylogeny: (Gorilla_gorilla_gorilla_western_lowland_gorilla:0.021599 ←)
9. Branch Number: 84 (Site, Probability, Transition) 42, 0.795, G>S 61, 0.823, T>S 119, 0.863, S>T 134, 0.875, S>P 148, 0.953, S>Q 149, 0.996, S>S Lineage in the phylogeny: (Callithrix_jacchus_whitetuftedear_marmoset:0.090719,((((Macaca_fascicularis_crabeating_macaque:0.006578,Macaca_mulatta_rh esus_monkey:0.000004):0.013118,Chlorocebus_sabaeus_green_monkey:0.019924):0.013720,(Rhinopithecus_bieti_black_snubno sed_monkey:0.000004,Rhinopithecus_roxellana_golden_snubnosed_monkey:0.000004):0.025800):0.006735,(Nomascus_leucogen ys_northern_whitecheeked_gibbon:0.051824,(Pongo_abelii_Sumatran_orangutan:0.021484,(Gorilla_gorilla_gorilla_western_lowlan d_gorilla:0.021599,(Homo_sapiens_human:0.000004,(Pan_paniscus_bonobo:0.000004,Pan_troglodytes_chimpanzee:0.000004):0. 013099):0.000004):0.041490):0.000004):0.017607):0.136167 ←)
10. Branch Number: 88 (Site, Probability, Transition) 43, 0.974, Q>P 78, 0.592, T>G 82, 0.697, G>S 103, 0.973, Q>P Lineage in the phylogeny: Loxodonta_africana_African_savanna_elephant:0.120966 ←

#### Predicted evolutionary rates for each branch of the phylogenetic tree (branch lengths are the dN/dS values predicted by CODEML)

((((((((((((((Zalophus_californianus #999.0000, Eumetopias_jubatus #244.3665) #225.7150, Odobenus_rosmarus #689.4730) #397.1745, (Leptonychotes_weddellii #0.0001, Monachus_monachus #949.6424) #999.0000) #0.1697, Enhydra_lutris #0.0001) #91.0856, Ursus_maritimus #3.2462) #0.0001, (Canis_lupus #999.0000, Vulpes_vulpes #0.5161) #0.2592) #51.8415, ((Panthera_tigris #116.6303, Panthera_pardus #999.0000) #263.3471, (Acinonyx_jubatus #638.3701, (Felis_catus #0.0001, Puma_concolor #293.8635) #336.2687) #271.5981) #0.2566) #0.0001, Manis_javanica #0.2341) #0.0001, (Ceratotherium_simum #0.2404, (Equus_asinus #0.0001, (Equus_caballus #195.2510, Equus_przewalskii #238.5228) #142.0174) #0.2435) #0.0001) #70.8301, (((Camelus_ferus #114.4290, Camelus_bactrianus #116.8703) #0.2461, Vicugna_pacos #0.0001) #0.2249, (Sus_scrofa #0.3767, ((((((Lagenorhynchus_obliquidens #0.0001, Orcinus_orca #999.0000) #0.0001, Delphinapterus_leucas #999.0000) #103.3377, Lipotes_vexillifer #0.5114) #0.0001, Physeter_catodon #0.2514) #999.0000, Balaenoptera_acutorostrata #999.0000) #0.6677, (((Bos_taurus #999.0000, Bison_bison #198.5425) #0.7618, Bubalus_bubalis #0.9989) #999.0000, (Pantholops_hodgsonii #63.5889, (Capra_hircus #171.4609, Ovis_aries #0.0001) #0.0001) #0.3264) #0.0001) #0.0001) #0.0001) #0.2646) #147.3301, (((Myotis_lucifugus #0.0001, Miniopterus_natalensis #0.0001) #144.6956, Desmodus_rotundus #0.7531) #1.3792, (Hipposideros_armiger #0.1710, (Rousettus_aegyptiacus #0.5025, (Pteropus_vampyrus #0.0001, Pteropus_alecto #71.8168) #0.0001) #0.1144) #27.6949) #1.1259) #0.0001, (Erinaceus_europaeus #0.0958, Condylura_cristata #0.2450) #1.2957) #0.0001, (((((((((Rattus_norvegicus #0.1228, Mus_musculus #0.0001) #0.0001, Meriones_unguiculatus #0.0001) #0.0001, (Peromyscus_maniculatus #0.0001, (Mesocricetus_auratus #0.0001, Cricetulus_griseus #0.0001) #0.0001) #135.4654) #0.0001, Nannospalax_galili #0.2119) #0.0001, Dipodomys_ordii #0.0001) #0.0001, (Urocitellus_parryii #183.2193, (Marmota_flaviventris #143.7105, Marmota_marmota #0.0001) #0.0001) #0.1417) #0.0001, (Heterocephalus_glaber #0.2017, (Cavia_porcellus #0.6168, Octodon_degus #0.2267) #0.0001) #0.0001) #0.0001, (Oryctolagus_cuniculus #0.1365, Ochotona_princeps #0.0358) #0.0001) #0.0001, (((Propithecus_coquereli #999.0000, Microcebus_murinus #0.0001) #0.0001, Otolemur_garnettii #0.0001) #999.0000, ((Aotus_nancymaae #0.3867, Cebus_capucinus #0.0001) #0.0001, (((((Pan_troglodytes #0.8493, Pan_paniscus #146.8477) #999.0000, Homo_sapiens #78.6724) #103.4384, Gorilla_gorilla #77.7181) #0.2426, Pongo_abelii #999.0000) #0.0001, (((Rhinopithecus_roxellana #128.4902, Rhinopithecus_bieti #0.9253) #0.4929, (Piliocolobus_tephrosceles #0.2836, Colobus_angolensis #95.7057) #102.5470) #87.3284, (Chlorocebus_sabaeus #0.0001, (((Macaca_mulatta #162.9342, Macaca_fascicularis #153.6423) #145.0405, Macaca_nemestrina #106.9026) #0.9058, ((Mandrillus_leucophaeus #73.4895, Cercocebus_atys #0.2822) #157.5496, (Papio_anubis #0.2842, Theropithecus_gelada #122.4933) #161.9117) #89.8523) #999.0000) #0.0001) #0.0001) #0.1470) #0.0001) #774.0332) #0.0001) #0.0001, (Dasypus_novemcinctus #0.2869, (Trichechus_manatus #0.3885, (Chrysochloris_asiatica #0.0889, Elephantulus_edwardii #0.0995) #0.4475) #0.0001) #0.0001) #0.0008, Gallus_gallus #0.0038);

#### Sequences of Q-rich and P-rich variants

>Q-rich SQSSSYGQPQSGSYSQQSSYGGQQQSYGQQQ

>P-rich PQSSSYGQPPSGSYSPQPSYGGQPPSYGQPP

